# A mean-field model of neural networks with PV and SOM interneurons reveals connectivity-based mechanisms of gamma oscillations

**DOI:** 10.1101/2025.10.23.684108

**Authors:** Farzin Tahvili, Martin Vinck, Matteo di Volo

**Author notes:** Equal contributing last authors (Martin Vinck, Matteo di Volo).

## Abstract

Classic theoretical models of cortical oscillations are based on the interactions between two populations of excitatory and inhibitory neurons. Nevertheless, experimental studies and network simulations suggest that interneuron subclasses such as parvalbumin (PV) and somatostatin (SOM) exert distinct control over oscillatory dynamics. Yet, we lack a theoretical understanding of the mechanisms underlying oscillations in E-PV-SOM circuits and of the differences with respect to the classical mechanisms for oscillations in simpler E–I networks. Here, we derive a biologically realistic mean-field model of a canonical three-population E-PV-SOM circuit. This model robustly generates oscillations whose features are consistent with experimental observations, including the relative timing of PV and SOM activity and the effects of optogenetic perturbations. By reducing the model to a linear analytical form, we demonstrate that gamma oscillations emerge directly from the cell-specific connectivity of the three-population circuit. This connectivity motif alone accounts for experimentally observed phase relationships, with PV activity consistently leading that of SOM neurons. Together, this mean field model identifies a distinct structural mechanism giving rise to oscillations in canonical E–PV–SOM circuits and provides theoretical primitives for constructing large-scale, cell-type-specific models of cortical dynamics.

## Introduction

Oscillations are a hallmark feature of neural dynamics, measurable from voltage membrane potentials to macroscopic electromagnetic waves, and are thought to play important roles in neural coding and signal transmission (Buzsáki, 2006; Wang et al., 2010; Vinck et al., 2023, 2025; Fries, 2005; Bressler and Kelso, 2001; Varela et al., 2001). Their prevalence in electrophysiological recordings is mirrored by their natural emergence in computational models of interacting excitatory and inhibitory spiking neurons, as well as in purely inhibitory spiking neuronal networks (Buzsáki et al., 2012; Brunel, 2000; Tiesinga and Sejnowski, 2009; Börgers and Kopell, 2003; Traub, 2006; Jadi and Sejnowski, 2014). However, while the theoretical underpinnings of oscillatory mechanisms in two-population models of excitatory and inhibitory neurons are well understood, we lack a theoretical understanding of cortical oscillations in more biologically realistic and heterogeneous microcircuits.

Mean-field models offer a powerful theoretical tool for dissecting the origin and stability of oscillations in neuronal networks. The pioneering Wilson–Cowan model showed that the population activity of Excitatory/Inhibitory (E/I) networks can be effectively captured by simple population equations, revealing the basic mechanisms underlying gamma oscillations (Wilson and Cowan, 1972). Thanks to its simplicity and generality, it has become a standard model for describing canonical mechanisms driving oscillatory regimes in E/I networks such as Pyramidal–Interneuron Gamma (PING) and Interneuron Gamma (ING) in purely inhibitory networks. Consistent with Wilson-Cowan models, inhibitory and excitatory neurons show distinct phase relationships to gamma oscillations in several cortical and hippocampal circuits (e.g., (Hasenstaub et al., 2005; Csicsvari et al., 2003; Buzsáki and Wang, 2012; Le Van Quyen et al., 2016; Onorato et al., 2025; Vinck et al., 2013; Onorato et al., 2020b; Cardin et al., 2009; Vinck et al., 2016)). Despite its elegance, the Wilson and Cowan model is a phenomenological model, as it is not derived directly from the spiking neural network it is supposed to represent. Over the years, considerable effort has been devoted to developing mean-field models capable of accurately reproducing the actual dynamics of spiking neural networks. In the context of quadratic integrate-and-fire (QIF) models, exact reductions have been derived to obtain low-dimensional representations of population dynamics (Montbrió et al., 2015). However, these models miss key biological features of neural networks that strongly affect population dynamics, such as sparse connectivity between neurons (Di Volo and Torcini, 2018) and employ *ad hoc* Cauchy distribution of heterogeneities or noise sources (Goldobin et al., 2021). Other mean-field approaches have instead aimed to increase biophysical realism, for instance by incorporating conductance-based synapses and spike-frequency adaptation in sparse spiking neural networks that mimic *in vivo* circuits (Di Volo et al., 2019). Regardless of the specific formulation, most of these models still rely on classical two-population E/I architectures, and thus on the PING or ING mechanisms proposed initially by the Wilson–Cowan framework.

Crucially, inhibitory neurons are not a homogeneous population but comprise distinct subtypes with specific circuit functions (Batista-Brito et al., 2018; Freund, 2003; Moore et al., 2010; Cardin, 2018; Miri et al., 2018). Among them, parvalbumin-expressing (PV) and somatostatin-expressing (SOM) interneurons have been shown to play complementary roles in controlling cortical oscillations (Veit et al., 2017; Chen et al., 2017a; Onorato et al., 2025). Strikingly, during gamma oscillations, the experimentally observed phase delay occurs relatively high in SOM cell spikes, whereas PV interneurons tend to oscillate nearly in phase with the excitatory population (Onorato et al., 2025). These experimental findings challenge the traditional view of gamma and beta oscillations as emerging from a single, generic E–I mechanism, thus calling for a revised theoretical framework that explicitly accounts for interneuron diversity. Several recent studies have employed neural network models of E, PV, and SOM cells to investigate the role of distinct interneuron subtypes in gamma oscillations (Lee et al., 2013; Moreni et al., 2025; Parker et al., 2025; Milea et al., 2025). Other works have instead used phenomenological neural mass models to show that distinct interneuron types exert different effects on population dynamics (Wendling et al., 2024; Hahn et al., 2022; Sanchez-Todo et al., 2023). In a recent work, we introduced the CAMINOS (Canonical Microcircuit Network Oscillations) model (Tahvili et al., 2025), a spiking neural network calibrated on experimental data, which proposed distinct roles of PV and SOM cells in generating and stabilizing gamma oscillations. While PV cells control mainly the frequency of oscillations, SOM cells control their amplitude. The CAMINOS model also was able to reproduce a broad range of experimental observations, in particular the effects of causal optogenetic manipulations targeting distinct interneuron classes during gamma oscillations. Numerical simulations revealed that a critical factor in the CAMINOS model was the specific asymmetric connectivity in canonical E-PV-SOM circuits, characterized by SOM cells exhibiting little to no self-inhibition and PV cells providing weak or no inhibition onto SOM cells.

However, because spiking network models are inherently high-dimensional and can only be explored through numerical simulations, it remains unclear whether the underlying computational mechanism in CAMINOS is distinct from a classical or effective PING model. Here, we take a crucial step by deriving a mean-field model of E–PV–SOM networks that preserves biological constraints on cellular properties and structural connectivity while remaining analytically tractable. We compared this mean-field model to direct numerical simulations of the corresponding spiking neural network and performed in silico manipulations on different interneuron populations within the mean-field framework - mimicking optogentic manipulations - to assess its capacity to predict experimental observations. We started with the biologically realistic mean-field model and progressively simplifyied the model to a linear skeleton, to analytically elucidate the existence of an oscillatory mechanism distinct from the classical PING model. Our results confirm that CAMINOS is a distinct oscillatory state emerging in three populations E-PV-SOM networks because of its special cell-specific connectivity structure.

## Results

The Results section is divided into two main parts. In the first part, we develop and analyze a biologically realistic mean-field model of the E–PV–SOM spiking network. We will investigate how this mean field model reproduces key experimentally observed features of cortical population dynamics. In the second part, we construct a linear mean-field model that preserves the essential circuit architecture of the biologically realistic model, but allows for an exact analytical treatment.

### Biologically realistic mean-field model of pyramidal, PV, and SOM populations

We constructed a spiking network model, consisting of three cell types: excitatory pyramidal cells (E cells), PV interneurons, and SOM interneurons. The network comprises 8,000 excitatory cells and 1,500 inhibitory interneurons, with half of the interneurons being PV and the other half being SOM. We considered the network to be random and sparse, with each neuron forming a synapse on another neuron with a probability of 0.07. We designed the connectivity structure between cell populations based on (Pfeffer et al., 2013), and we point out two key features: the absence of PV-to-SOM inhibition and the absence of self-inhibition of SOM. The dynamics of each neuron were modeled using the Adaptive exponential Integrate-and-fire (AdEx) model (Brette and Gerstner, 2005), and the couplings between neurons were modeled as conductance-based synapses. A synaptic delay of 2 ms was included for all synapses. We refer the reader to the Methods section, where detailed explanations, equations, and parameter values are provided. The schematic of the designed canonical microcircuit is shown in Fig. 1A.

**Figure 1.**
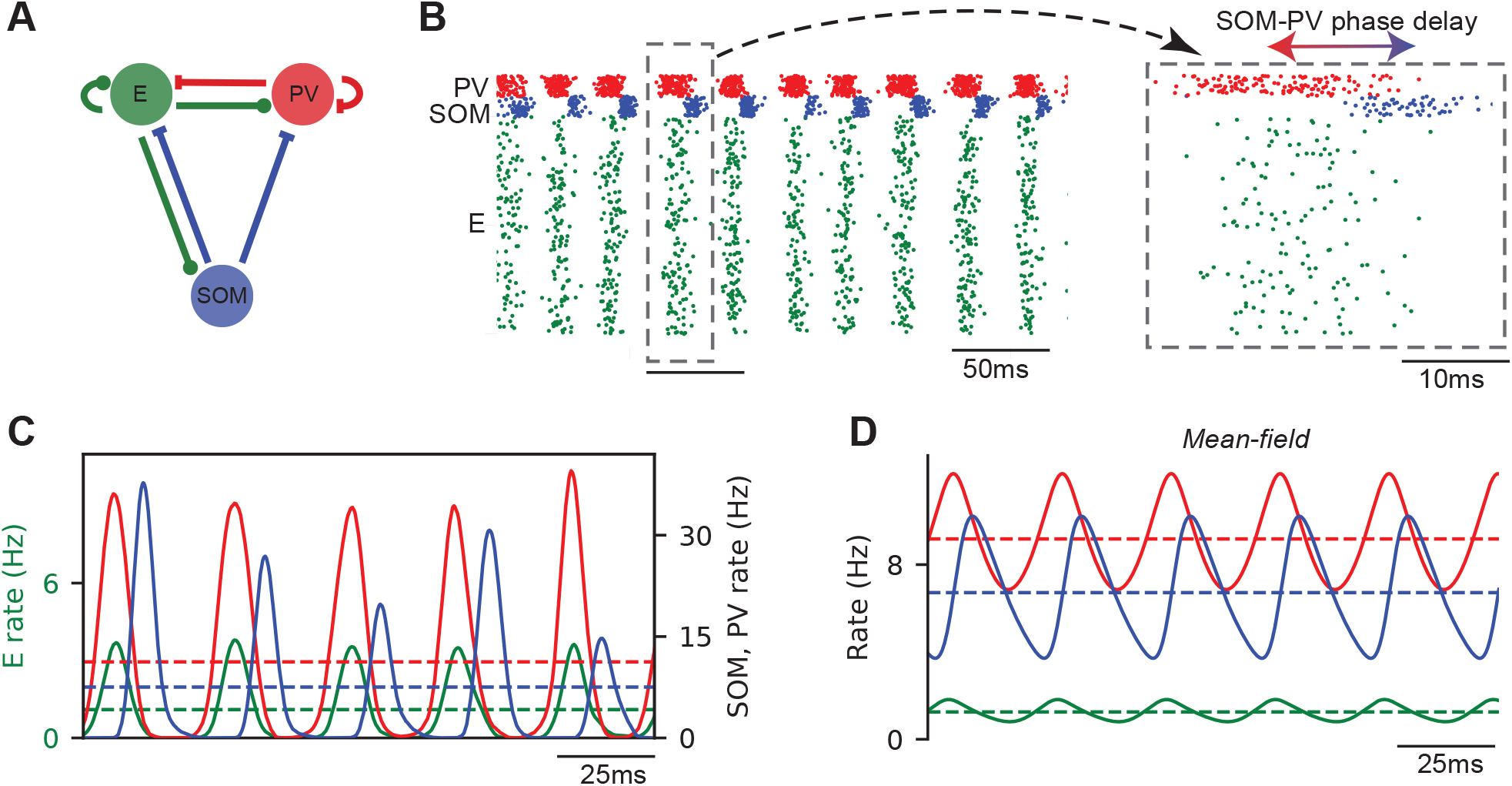
Biologically realistic mean field model setup and emergent dynamics. (A) Schematic of the E–PV–SOM circuit. (B) Raster plot of the simulated spiking network, with a zoomed-in view of one gamma cycle. (C) Population activity traces of the spiking network. (D) Corresponding mean-field simulation. In all panels, green represents E neurons, red represents PV interneurons, and blue represents SOM interneurons.

The network naturally generates gamma oscillations, as can be observed from the raster plot shown in Fig. 1B. The emergent dynamics stabilizes in an oscillatory regime with an oscillation frequency of approximately 35 *Hz* within the low-gamma range, consistent with experimental results (Onorato et al., 2025; Veit et al., 2017). Fig. 1B illustrates that SOM cells consistently fire with some delay relative to PV cells. This phase delay was estimated quantitatively across gamma cycles using the approach described in (Tahvili et al., 2025) and is found to be around 8.5 ms. The population activities of the spiking network are shown in Fig. 1C, which illustrates that PV cells exhibit the highest mean firing rate, followed by SOM interneurons, and finally E cells with the lowest mean firing rate. We note that the mean firing rates, oscillation frequency, and phase delay between PV and SOM interneurons agree with previous experimental findings (Onorato et al., 2025; Veit et al., 2017; Spyropoulos et al., 2022).

The next step was to derive a mean-field approximation and compare it to the spiking network. We used the mean field modeling approach developed for AdEx neurons in (Di Volo et al., 2019). This approach basically employs a master equation formalism first developed in (El Boustani et al., 2009), that assumes the network in an asynchronous irregular state, memoryless over a time scale *T*. At the first order, the equations are formally the same as in the Wilson–Cowan model, but instead of a sigmoid, neuron’s firing activity is driven by biophysically grounded transfer functions. These transfer functions are derived from fluctuation-driven single-neuron dynamics, following the approach of (Zerlaut et al., 2016) that we briefly summerise hereafter. In vivo–like cortical neurons, as neurons in our spiking network, operate in a high-conductance state (Destexhe et al., 2003) characterized by large synaptic conductances. This regime arises from dense, irregular poissonian synaptic bombardment, which produces stochastic fluctuations in the membrane potential around a depolarized mean near a spiking threshold. From these fluctuations, one can construct a transfer function that deterministically maps the statistical properties of the membrane voltage—its mean (*µ*_*v*_), variance 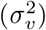, and effective time constant (*τ*_*v*_)—to the expected firing rate of a neuron. By expressing these membrane voltage statistics as functions of pre-synaptic population rates, the transfer function provides a principled input–output mapping for the population dynamics. One should note that generally 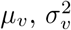, and *τ*_*v*_ are not constants and evolve through time. We refer the reader to the Methods section, where we provide a detailed explanation of the derivation of transfer functions.

As we considered AdEx neurons we also introduced an adaptation variable *w* for each neuronal population in our mean-field equations, representing the average adaptation current within each group, see Di Volo et al. (2019). The two key parameters, *a* (subthreshold adaptation) and *b* (spike-triggered adaptation), control the contribution of membrane voltage fluctuations and spike events to this current, respectively. The adaptation time scale, denoted by *τ*_*w*_, is assumed to be much slower than the mean-field time scale *T* and therefore, on the mean-field timescale, the adaptation variable *w* can be treated as stationary, allowing the population firing rates to be computed conditionally on the current value of *w* while the slow adaptation dynamics evolve over a longer timescale. The firing rates determine the evolution of adaptation, while adaptation in turn modifies the membrane voltage statistics (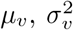, and *τ*_*v*_) and consequently affects the transfer function. In this way, the mean-field equations capture both fast fluctuation-driven dynamics and slow negative feedback from adaptation. The timescale of adaptation is the same as that of the spiking network (500 ms; see Methods section). The time constant *T* of the mean-field model is of the same order of the membrane time constant, and it was here chosen to be 15 ms such that the resulting dynamics reproduced the network’s temporal behavior and oscillation frequency. Finally, synaptic delay is introduced following the approach developed in Tahvili and Destexhe (2024).

The mean-field model is then described by the following set of equations 1 in which *ν*^*∗*^ represents the population firing rates of the cell type *∗* ∈ *{E, PV, SOM }*. ℱ _*∗*_ shows the transfer functions, 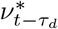 denotes *ν* delayed by *τ*_d_ = 2 *ms* representing the synaptic delay. 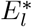 represents the resting membrane potential and *a*_*∗*_ and *b*_*∗*_ are the adaptation parameters with the same values as for the spiking network. It should be noted that the parameters of the spiking network, such as connection probabilities, number of neurons, synaptic properties, and neurons’ biophysical parameters, were incorporated in the derivation of the transfer functions. In other words, the transfer functions were calculated and fitted for specific values of the parameters of the spiking network. We refer the reader to the Methods section, where we provide a detailed explanation of the derivation of transfer functions based on the spiking network.

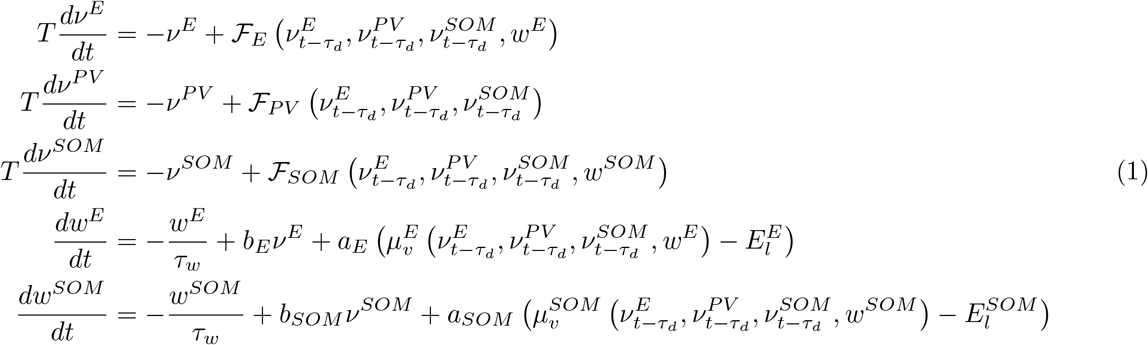

Fig. 1D shows the mean-field simulation, which produces a robust oscillatory regime at 35 *Hz*, with a PV–SOM phase delay of approximately 7 *ms*. We have found good agreement between the mean-field and spiking network models in key proprieties, such as mean firing rates, frequency of oscillation, and PV–SOM phase delay. It is important to notice that the peak of activity of SOM and PV is higher in the network simulations with respect to the mean field model. It is natural to expect some quantitative discrepancies in this model, given the approximations performed in the transfer function evaluation. Nevertheless, the agreement with a sparse biophysical networks is overall very good in terms of dynamical behavior and key quantities in relation to experimental data (mean rates, oscillation frequency and neurons’ spike timing).

Throughout the study, we define the mean-field (and the corresponding spiking network) described in this section as the “reference model”. In subsequent sections, when we refer to the reference model, we mean this original setup.

### Distinct contributions of PV and SOM interneurons to gamma oscillations

In experiments, optogenetic silencing has been used to investigate the causal contributions of specific interneuron classes to cortical dynamics (Veit et al., 2017; Chen et al., 2017a). We dissected the contributions of PV and SOM interneurons in our oscillation-generating model using two types of analyses, which we term “silencing” and “randomizing.” By silencing a given proportion of a cell type, we mean reducing the number of neurons of that type by the same proportion in the reference model. By randomizing a given proportion of a cell type, we mean randomizing the spike times of that proportion—that is, replacing that fraction of neurons with Poisson units emitting spikes at a constant rate equal to the mean population activity of that cell type in the reference model.

The top panel of Fig. 2A shows the mean-field dynamics for PV silencing levels ranging from 0% to 100%. Increasing the proportion of silenced PV neurons results in elevated firing rates and larger oscillation amplitudes, accompanied by a reduction in oscillation frequency. When the fraction of silenced PV neurons exceeds 70%, the system undergoes a phase transition into a seizure-like regime characterized by the maximum firing rate of the excitatory population, equal to the inverse of the refractory period. These findings are in agrreement with direct numerical simulations of the spiking neural networks. Representative raster plots and mean-field traces for 40% and 90% PV silencing are shown in Fig. 2A.

**Figure 2.**
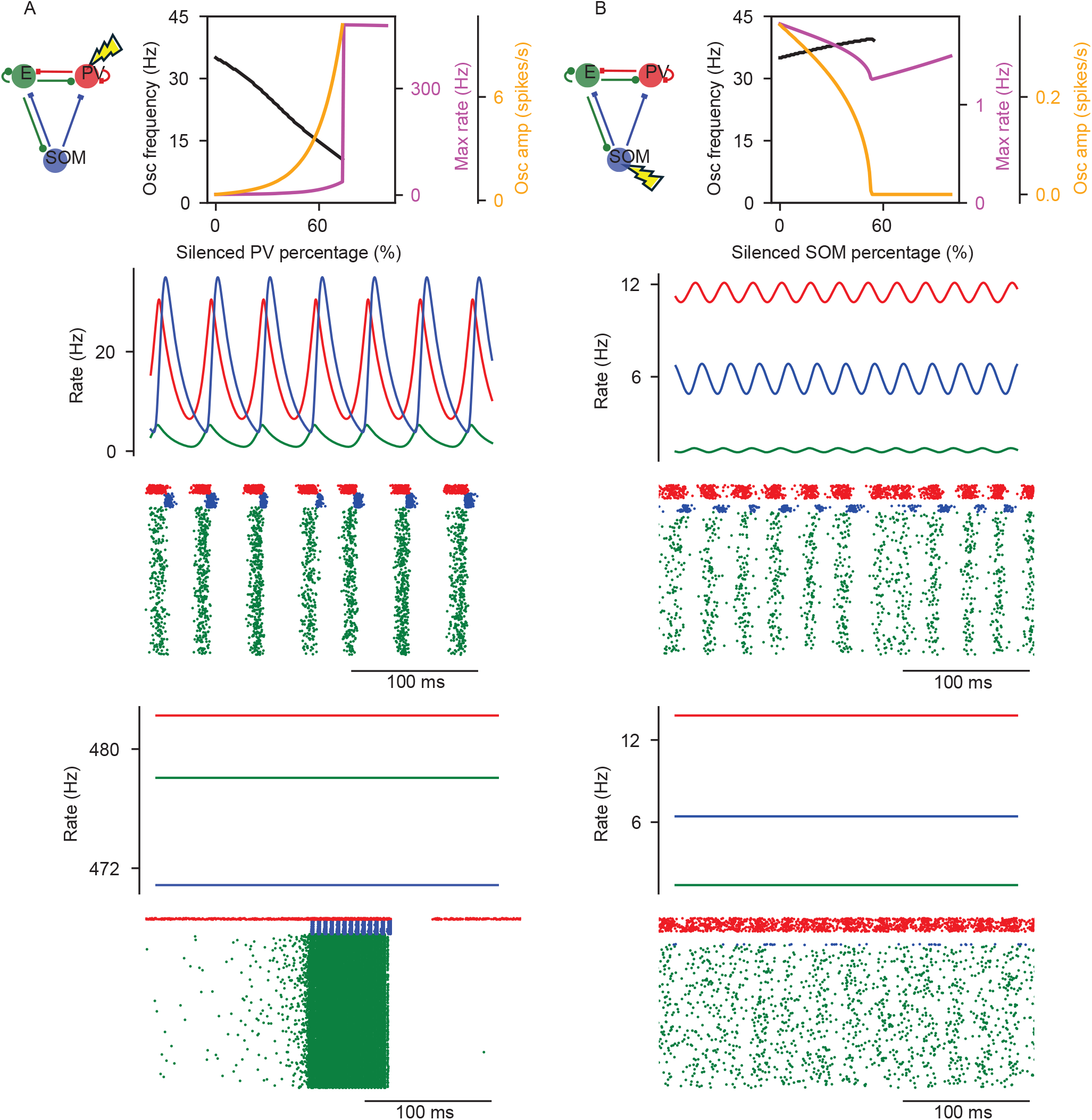
Emergent dynamics across different levels of interneuron silencing. Column (A) shows the results of silencing PV interneurons, and column (B) shows the corresponding results for silencing SOM interneurons. In column (A), the top panel depicts the mean-field characteristics across different levels of PV silencing: the black line indicates oscillation frequency, the orange line indicates oscillation amplitude, and the magenta line shows the maximum firing rate of the excitatory population. The two middle panels present the spiking network raster plot and mean-field activity when 40% of PV interneurons were silenced. The bottom panels show the raster plot and mean-field activity when 90% of PV interneurons were silenced. Column (B) follows the same structure but for silencing SOM interneurons. In all panels, green represents excitatory neurons, red represents PV interneurons, and blue represents SOM interneurons.

Randomization of PV neurons produced qualitatively similar outcomes, with minor variations (Fig. 3A, top panel). Representative raster plots and mean-field traces for 40% and 90% PV randomization are also shown in Fig. 3A. Overall, these findings—consistent with experimental observations (Veit et al., 2017) underscore the fundamental role of PV cells in maintaining circuit stability and further suggest their critical contribution to regulating both the frequency and amplitude of oscillations.

**Figure 3.**
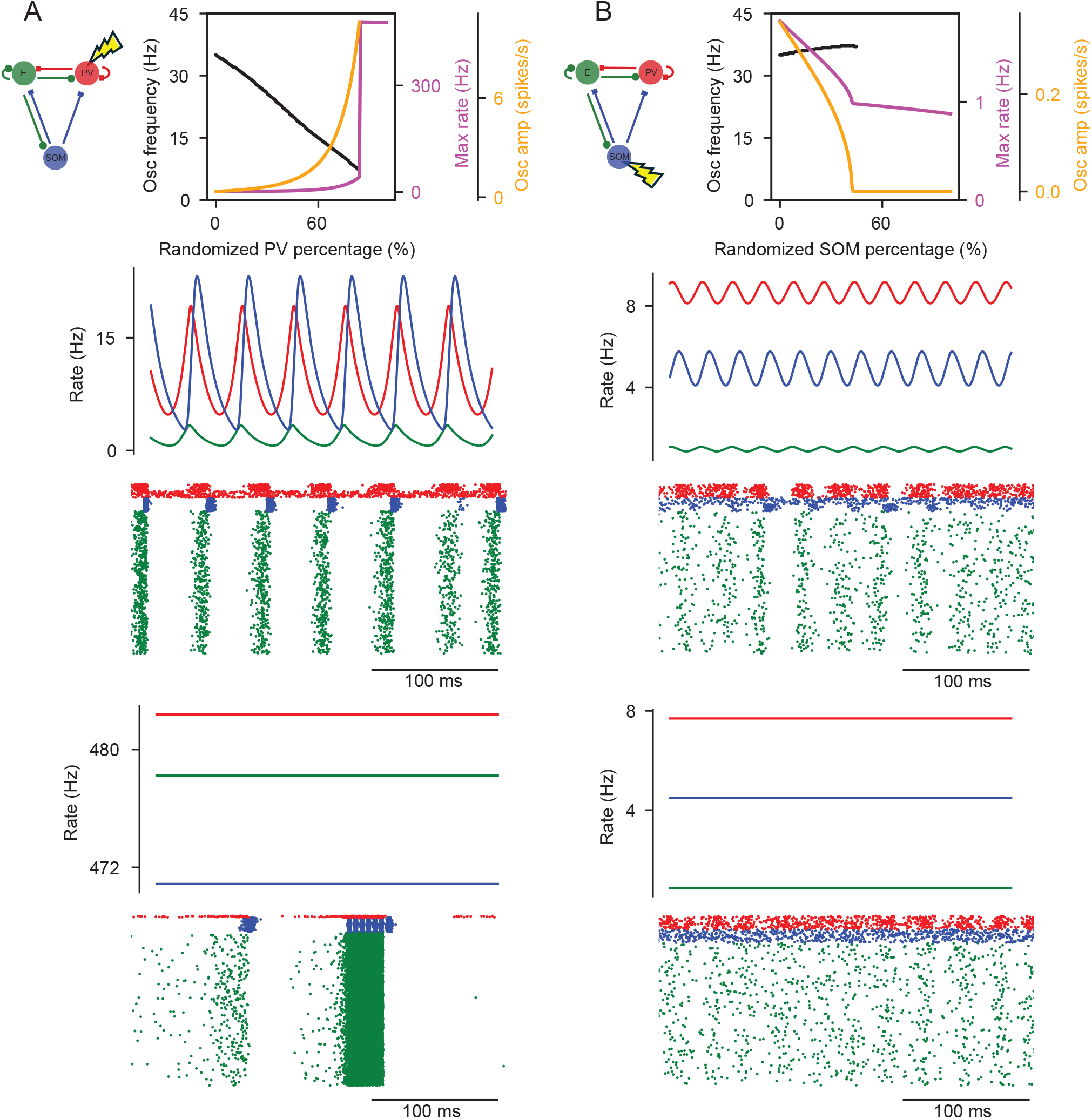
Emergent dynamics across different levels of interneuron spike times’ randomization. Column (A) shows the results for randomizing PV interneuron spike times, and column (B) shows the corresponding results for SOM interneurons. In column (A), the top panel shows the mean-field dynamics across different levels of PV spike-time randomization: the black line indicates oscillation frequency, the orange line indicates oscillation amplitude, and the magenta line shows the maximum firing rate of the excitatory population. The two middle panels present the spiking network raster plot and mean-field activity when 40% of PV spikes are randomized. The bottom panels show the raster plot and mean-field activity when 90% of PV spikes are randomized. Column (B) follows the same structure, but for randomizing SOM interneuron spikes. In all panels, green represents excitatory neurons, red represents PV interneurons, and blue represents SOM interneurons.

The top panel of Fig. 2B shows the mean-field dynamics under different levels of SOM silencing. Increasing the proportion of silenced SOM neurons leads to a progressive reduction in oscillation amplitude, and at 60% silencing, the system undergoes a bifurcation, transitioning into a stable non-oscillatory regime. The oscillation frequency also increases slightly with SOM silencing, although this effect was much smaller than the frequency shift observed during PV silencing. Representative raster plots and mean-field traces for 40% and 90% SOM silencing are shown in Fig. 2B. These findings recapitulate previous experimental observations (Veit et al., 2017; Chen et al., 2017a).

Randomization of SOM neurons produced qualitatively similar outcomes, with minor variations (top panel of Fig. 3B). Representative raster plots and mean-field traces for 40% and 90% SOM randomization are also shown in Fig. 3B. Overall, these results, consistent with experimental findings, demonstrate that SOM interneurons, and their spike timing, play a key role in sustaining oscillatory amplitude, as their silencing and or randomizing abolishes rhythmic activity.

### Gradients in interneuron densities shape oscillatory regimes and vulnerability to epilepsy

PV and SOM interneurons exhibit distinct laminar and regional distributions. PV cells reach their peak density in cortical layer 4, whereas SOM interneurons are most abundant in layer 5 (Tremblay et al., 2016; Kim et al., 2017). Moreover, the density ratio of SOM to PV interneurons varies systematically across cortical regions, forming a gradient from early sensory areas to higher-order association cortices as shown in the top panel of Fig. 4A (Murray et al., 2014). In primary sensory regions, the SOM/PV ratio is relatively low, indicating a predominance of PV interneurons, whereas in higher-order regions such as the PFC, the ratio is much higher, reflecting the great prevalence of SOM cells. Likewise, there is evidence that faster oscillations (gamma) are more prevalent in lower hierarchical levels and superficial and granular layers, while slower oscillations are more prevalent in higher hierarchical levels and deeper layers (Vinck et al., 2025; Bastos et al., 2015; Vezoli et al., 2020, 2021; Murray et al., 2014; Honey et al., 2012; Buffalo et al., 2011; Vinck et al., 2023).

**Figure 4.**
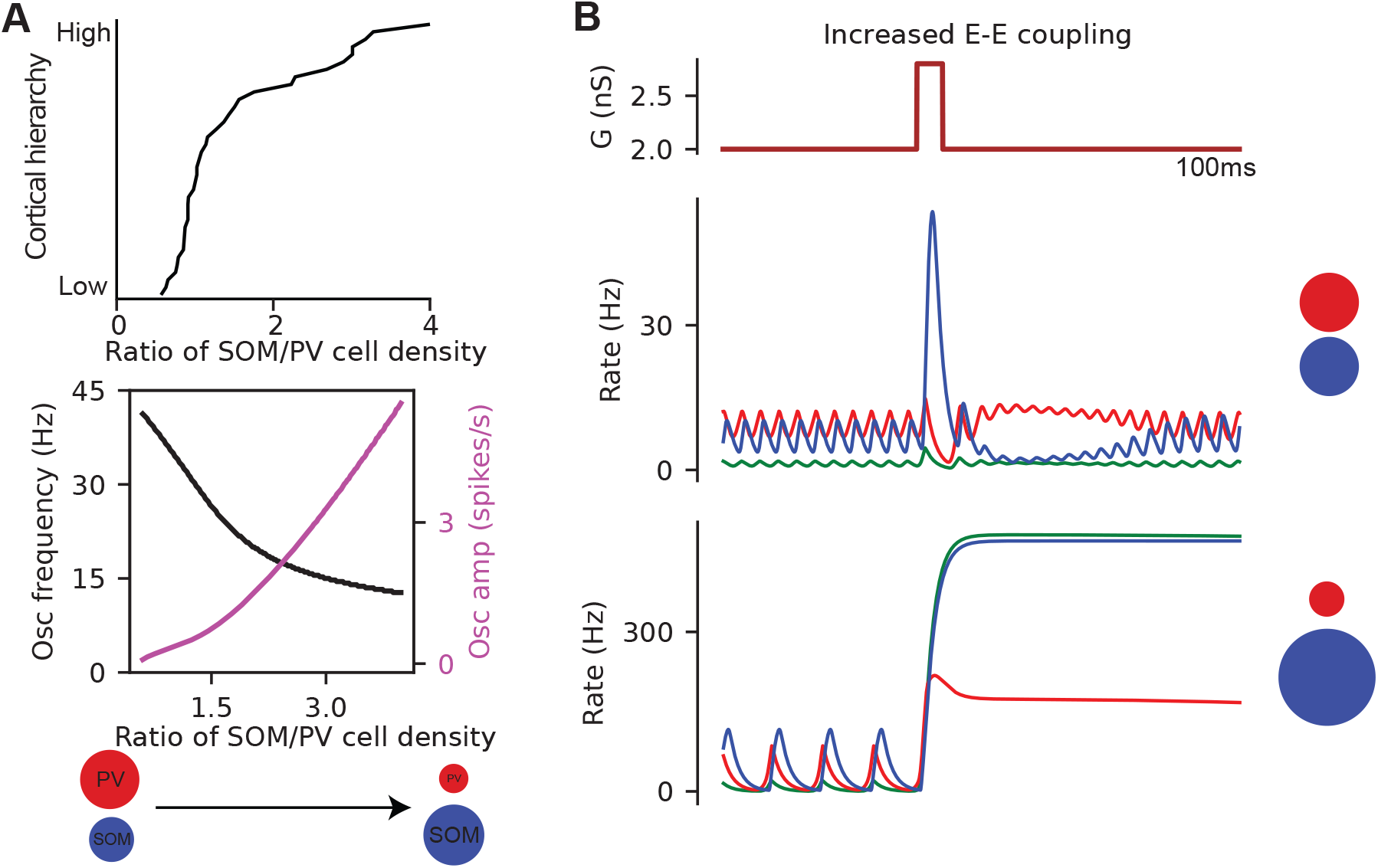
Influence of SOM/PV density ratio gradients on the emergent mean field dynamics. (A) Top panel: density ratio of SOM to PV interneurons across the cortical hierarchy, extracted from (Murray et al., 2014). (A) Bottom panel: meanfield dynamics at different SOM/PV density ratios, with the black line indicating oscillation frequency and the magenta line indicating oscillation amplitude. (B) Effect of a transient increase in excitatory self-coupling. The top panel of column (B) shows the applied increase transient in excitatory self-coupling. The middle panel presents the mean-field response to this transient when *SOM/PV* = 1, and the bottom panel shows the corresponding response when *SOM/PV* = 4.

Our model directly links oscillation frequency to the density of SOM and PV interneurons across cortical layers and regions. When the SOM/PV density ratio is high, the model generates slower oscillations. Conversely, when the SOM/PV density ratio is low, as in upper layers and/or sensory/motor cortices, the model produces faster oscillations. This correspondence demonstrates that variations in interneuron composition can account for the laminar and regional patterns of oscillatory activity observed experimentally. The mean-field dynamics under different SOM/PV density ratios are shown in Fig. 4A (bottom panel). As illustrated, increasing the SOM/PV ratio decreases the oscillation frequency while increasing its amplitude. Notably, when varying the SOM/PV density ratio, the total number of interneurons was kept fixed at 1500, with only the relative numbers of SOM and PV cells adjusted.

Next, we asked whether the balance between SOM and PV interneurons influences the stability of dynamics. To test this, we introduced a transient increase in excitatory-to-excitatory (E–E) coupling. As shown in Fig. 4B, the setup with a high SOM/PV ratio was more susceptible to instability under this perturbation, exhibiting seizure-like dynamics marked by a phase transition. In contrast, set-ups with a low SOM/PV ratio remained stable when the recurrent excitation transiently increased.

### Connectivity architecture, not synaptic kinetics or adaptation, as the fundamental mechanism of oscillation generation

Firstly, we investigate how the specific connectivity patterns contribute to the emergence of oscillatory activity and ask whether the particular arrangement of connections between cell types underlies the generation of oscillatory regimes. The key aspect of the reference circuit is that the SOM population neither inhibits itself nor receives inhibition from the PV population. To assess the functional relevance of this wiring scheme, we systematically introduce these missing connections one at a time and examine their individual impact on network dynamics.

Fig. 5A illustrates the effect of introducing additional inhibitory connections into the reference model. When a coupling from PV to SOM is added, increasing the strength of this coupling progressively reduces the amplitude of oscillation until a critical point is reached, at which a phase transition occurs and the system shifts into a non-oscillatory, stable regime. A similar effect is observed when self-inhibition is introduced onto the SOM population: strengthening this coupling also diminishes oscillation amplitude and eventually drives the system into a non-oscillatory stable state. However, the impact of the PV→SOM connection is much more pronounced, as the transition to the non-oscillatory regime occurs at lower connectivity strengths compared to the case of SOM self-inhibition. It should be noted that these additional inhibitory couplings (PV→SOM and SOM→SOM) were implemented within the same network framework described previously. As with the other connections, they were introduced as sparse, random synapses with the same connection probability (7%) and modeled using conductance-based synapses with identical synaptic delays of 2 *ms*.

**Figure 5.**
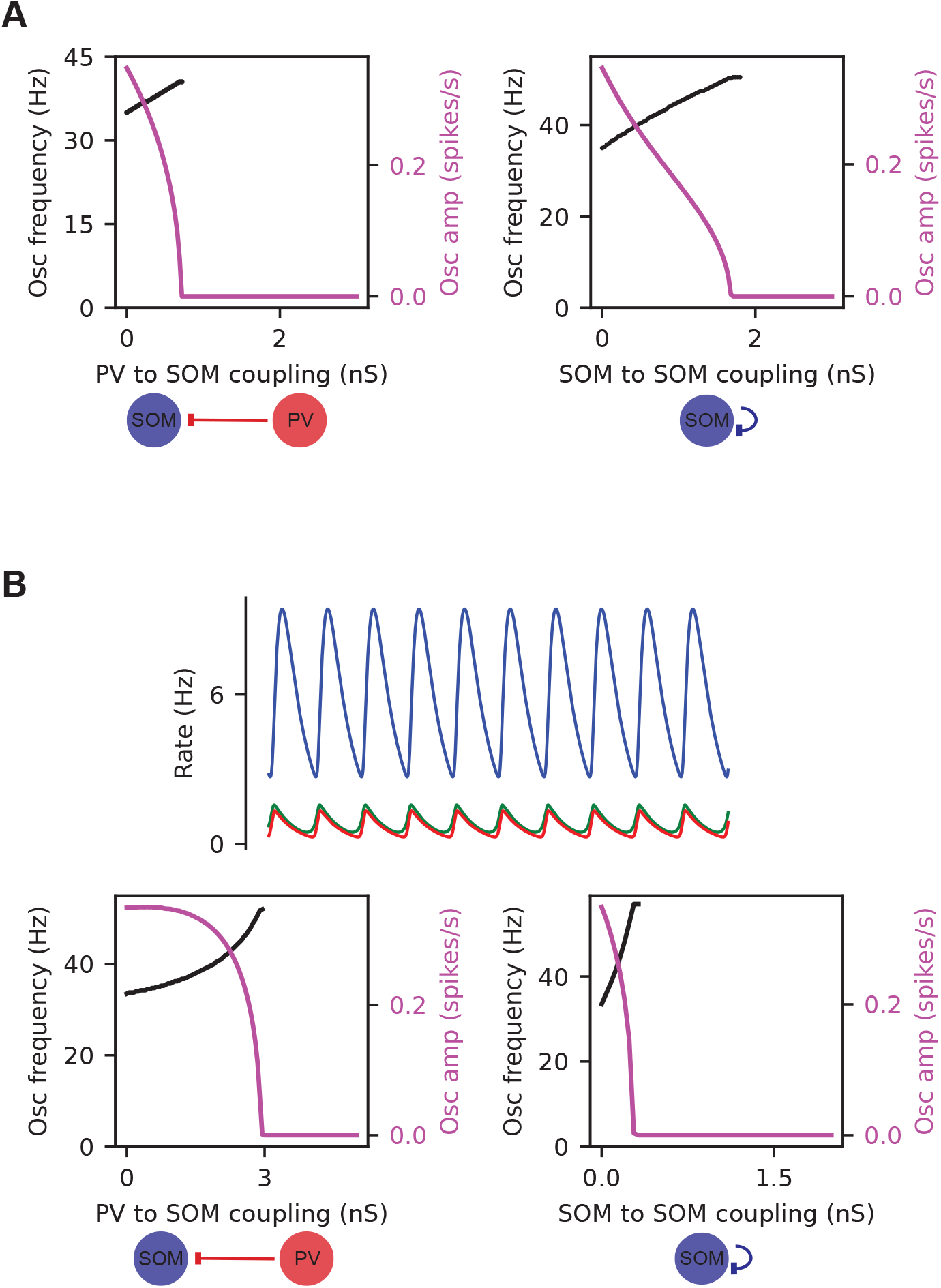
Role of SOM to SOM and PV to SOM connections. (A) Left: effect of PV→SOM inhibition on the mean-field dynamics. Right: effect of SOM self-inhibition on the mean-field dynamics. (B) Results for the reduced system (without synaptic delay and adaptation). Top: corresponding mean-field dynamics. Bottom: effects of PV→SOM inhibition (left) and SOM self-inhibition (right) on the reduced mean-field dynamics.

We next asked whether the fundamental mechanism underlying oscillatory dynamics lies solely in the connectivity architecture of the circuit, with other factors such as synaptic time constants/delay and spike frequency adaptation serving only as modulators rather than essential components. To test this hypothesis, we removed both synaptic timescales and adaptation from the model. In this simplified configuration, we also set the SOM leak reversal potential to 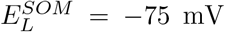 (to prevent the SOM overactivity) and provided external input only to the excitatory population, with a rate of 1 Hz. After removing synaptic timescales and adaptation, the mean-field equations take the following form:

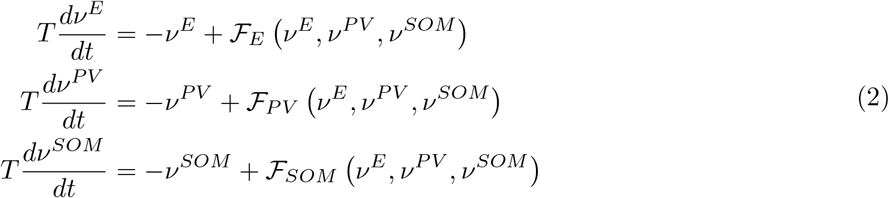

The mean-field equations 2 represent the version of the reference model without synaptic timescales/delay and adaptation. The upper panel of Fig. 5B illustrates the simulation of 2. In particular, even in the absence of synaptic and adaptation timescales, the system continues to generate oscillations in the low gamma range (the oscillation frequency is around 35 Hz). Moreover, PV activity consistently leads SOM activity in phase. These results demonstrate that the core mechanism driving this type of oscillation does not rely on synaptic timescales or the biophysical properties of neurons, such as adaptation. Instead, it suggests that the essential factor is the specific connectivity architecture—namely, the minimal or absent PV→SOM inhibition and the weak or absent SOM self-inhibition.

To further verify our findings, we reintroduced the PV→SOM and SOM→SOM connections separately into the reduced mean-field 2 and examined their effects on the dynamics. The lower left panel of Fig. 5B shows the results for the PV→SOM case: as the strength of this connection increases, the amplitude of oscillations progressively declines until a phase transition occurs, driving the system into a non-oscillatory stable regime. Similarly, the lower right panel of Fig. 5B illustrates the effects of adding self-inhibition to the SOM. Strengthening this connection likewise reduces the oscillation amplitude, and beyond a critical point, the system again transitions into a non-oscillatory stable state.

In summary, these analyses demonstrate that the essential driver of the oscillatory regime is the specific connectivity architecture, characterized by little to no PV→SOM inhibition and minimal SOM self-inhibition, rather than synaptic or adaptation timescales. This emphasizes that the structural wiring of the network, rather than its biophysical details, forms the fundamental basis of the oscillatory dynamics.

## Linear mean-field model

Assuming that a general equilibrium exists in the state space of a biological circuit composed of three cell types, one can reasonably expect that the system behaves linearly in the vicinity of this equilibrium. A linear approximation is then often valid for describing the dynamics around the equilibrium. On this basis, we develop and analyze a linear model of the E-PV-SOM circuit, demonstrating that some important features of the original system are retained in this simplified linear framework. This part presents a fully analytical development of the linear model; however, to keep the exposition accessible to a wide readership, we include the detailed mathematical proofs and technical arguments in the Appendix.

### Framework for the general linear rate model

We model the dynamics of population activity in a local cortical microcircuit comprising excitatory (E) neurons, parvalbumin-expressing (PV) interneurons, and somatostatin-expressing (SOM) interneurons. The population activity vector is defined as

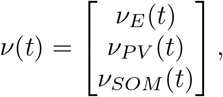

where each component represents the average firing rate of the corresponding neuronal population at time *t*. The temporal evolution of these activities is described by the linear rate model:

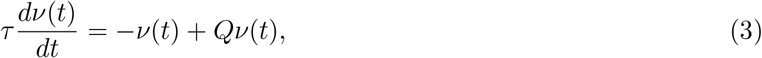

where *τ* denotes the time constant of population activity, and *Q* is the effective connectivity matrix:

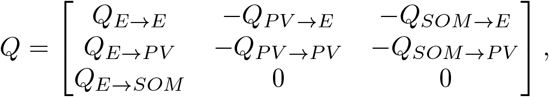

with *Q*_*∗*_ *>* 0 representing the effective coupling strength between populations. Positive entries correspond to excitatory projections originating from E neurons, while negative entries represent inhibitory influences mediated by PV and SOM interneurons. In this abstraction, we omit the inhibitory connections *PV → SOM* and *SOM → SOM*, consistent with experimental findings indicating that such synapses are very sparse or functionally negligible in cortical networks.

The term −*ν*(*t*) in Eq. 3 describes the intrinsic relaxation of population activity toward baseline in the absence of input. Biologically, this captures processes such as membrane leakage or adaptation that cause neural activity to decay when it is not sustained by excitation.

Defining *A* = *Q* − *I*, where *I* denotes the identity matrix, the system can be rewritten in the standard linear form:

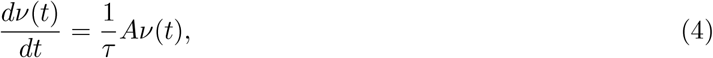

with

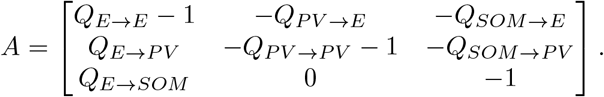

This formulation highlights the interplay between recurrent excitation and inhibition in shaping the temporal evolution of population activity. In general, the solution to Eq. 4 can be expressed through the eigen-decomposition of *A*:

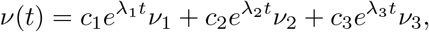

where *λ*_1_, *λ*_2_, *λ*_3_ are the eigenvalues of 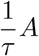 and *ν*_1_, *ν*_2_, *ν*_3_ are the corresponding eigenvectors. The constants *c*_1_, *c*_2_, *c*_3_ are uniquely determined by the initial population activity *ν*(0).

Each eigenmode (*λ*_*i*_, *ν*_*i*_) represents a distinct dynamical motif of the circuit, corresponding to coordinated patterns of population activity. The nature of the eigenvalues determines whether these modes decay, oscillate, or grow over time, thereby linking the circuit’s connectivity structure to its emergent temporal dynamics. We first clarify some terminology that we use regarding oscillatory dynamical regimes. Generally, oscillatory dynamics occur when the matrix *A* has a pair of complex conjugate eigenvalues and a negative (≤ 0) real eigenvalue. The damped oscillation arises when the complex conjugate pair has a strictly negative real part. The pure oscillation occurs when the complex conjugate pair is purely imaginary. The growing oscillation arises when the complex conjugate has a strictly positive real part.

### If E-PV-SOM circuit oscillates, then E and PV activity lead SOM activity in phase

In the linear system Eq. 4, oscillatory dynamics arises when the matrix *A* has complex-conjugate eigenvalues. Specifically, if *A* possesses an eigenvalue of the form *α* + *iω* with *ω >* 0, the corresponding eigenmode produces oscillations at frequency *ω/*(2*πτ*).

One of the key dynamical features of gamma oscillations in the E-PV-SOM circuit, as noted earlier, is that PV interneurons activity leads SOM cells activity in phase. To represent this phenomenon within the linear system, we note that if

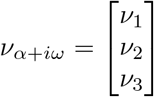

is an eigenvector of *A* associated with the eigenvalue *α* + *iω* (*ω >* 0), then the corresponding oscillatory mode exhibits a well-defined phase relationship among its components. In particular, a phase lead of the PV population relative to the SOM population is expressed by

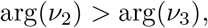

or equivalently, if 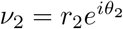 and 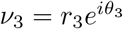, then *θ*_2_ *> θ*_3_.

In the following proposition, we prove that for any set of parameters *Q*_*∗*_ *>* 0, if *A* admits a non-real eigenvalue *α* + *iω* (*ω >* 0) with corresponding eigenvector *ν* = [*ν*_1_, *ν*_2_, *ν*_3_]^⊤^, the phases of *ν*_1_ (E population) and *ν*_2_ (PV population) are strictly larger than that of *ν*_3_ (SOM population). In other words, whenever Eq. 3 exhibits oscillatory behavior, the E and PV populations necessarily lead the SOM population in phase. We present the proposition here; the full mathematical proof, including technical details, is given in Appendix A.

#### Proposition 1.

*Suppose that the matrix*

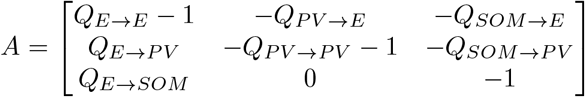

*in which all Q*_*∗*_ *are strictly positive, has a non-real complex eigenvalue α* + *iω and ν* = [*ν*_1_, *ν*_2_, *ν*_3_]^*T*^ *is the corresponding eigenvector. Then the phases of ν*_1_ *and ν*_2_ *are strictly larger than the phase of ν*_3_.

### Conditions for sustained oscillations in the E–PV–SOM linear system

In this section, we determine the conditions that must be maintained so that the solution of this system is purely oscillatory (not growing or damped oscillations). Having such a solution implies that 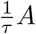 has a pair of pure imaginary *±iω*, (*ω >* 0) and a real *r* ≤ 0 eigenvalues. The characteristic polynomial of 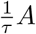 is

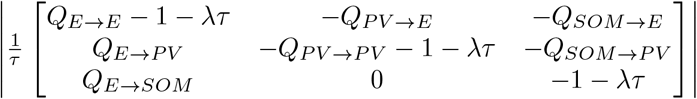

and must be equal to (*λ* + *r*)(*λ*^2^ + *ω*^2^). In other words, det 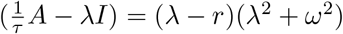 is required.

Expanding both sides and then matching the coefficients, yields the necessary and sufficient conditions as follows.

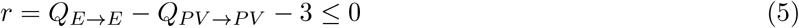

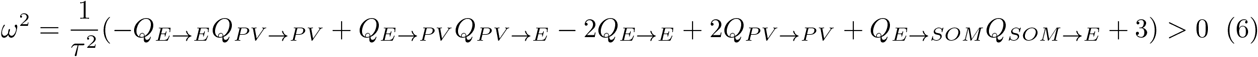

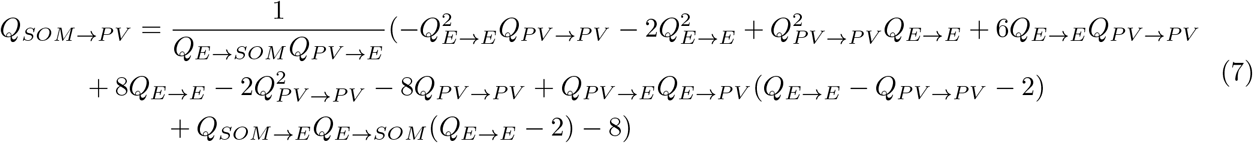

These three conditions also give us some intuitive insights into the system. For example, from 5, we understand that self-excitation cannot be much larger than PV self-inhibition. Equation 6, which is also an explicit expression of oscillation frequency 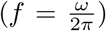, indicates that the oscillation frequency is mainly determined by the time constant *τ* since it depends linearly on 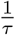 whereas it depends on the square root of *Q*_*∗*_. The oscillation frequency is also an increasing function of *Q*_*SOM*→*E*_ and *Q*_*E*→*SOM*_. Another property is that the oscillation frequency does not depend on *Q*_*SOM*→*PV*_.

### Parameter selection and dynamical regimes

The next stage is to assign values to parameters in order to specify our model. To this end, we first consider a system consisting solely of E and PV, assign values to its parameters to generate a damped oscillation, and demonstrate how adding SOM to this system sustains the oscillation. We consider that the system is 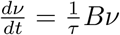 where

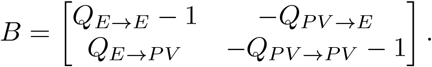

To have a damped oscillation in the E-PV system, the eigenvalues of *B* must be *γ ± iω*^*′*^ such that *γ <* 0, *ω*^*′*^ *>* 0. For these eigenvalues to exist, the following two conditions must be met.

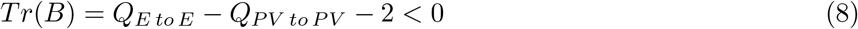

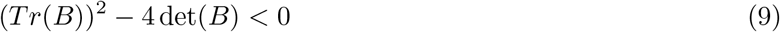

Assigning values to *Q*_*E*→*E*_, *Q*_*E*→*PV*_, *Q*_*PV* →*E*_, and *Q*_*PV* →*PV*_ must be such that it satisfies 8 and 9. We take *Q*_*E*→*E*_ = 5, *Q*_*E*→*PV*_ = 5, *Q*_*PV* →*E*_ = 5 and *Q*_*PV* →*PV*_ = 4.

To incorporate SOM into the system, it is important to note that satisfying 8 and 9 automatically ensures that 5 and 6 are also satisfied. Therefore, any choice of values for *Q*_*E*→*SOM*_, *Q*_*SOM*→*E*_, and *Q*_*SOM*→*PV*_ will inherently meet 5 and 6. Consequently, when selecting *Q*_*E*→*SOM*_, *Q*_*SOM*→*E*_, and *Q*_*SOM*→*PV*_, only 7 must be imposed.

We choose *Q*_*E*→*E*_ = 5, *Q*_*E*→*PV*_ = 5, *Q*_*PV* →*E*_ = 5, *Q*_*PV* →*PV*_ = 4, *Q*_*E*→*SOM*_ = 4. Then 7 gives 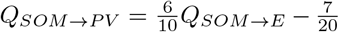. We choose *Q*_*SOM*→*E*_ = 4 and so *Q*_*SOM*→*PV*_ = 2.05. One should note that choosing these values for *Q*_*∗*_ is only an example. In addition, we take *τ* to be 20 *ms*. Therefore, our base linear model is:

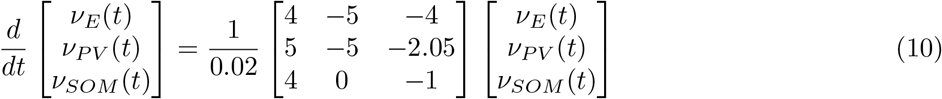

One can see the activity dynamics of the base linear model 10 in Figure 6A. The frequency of oscillation is around 40 *Hz*, and PV has a 4 *ms* phase advance relative to SOM.

**Figure 6.**
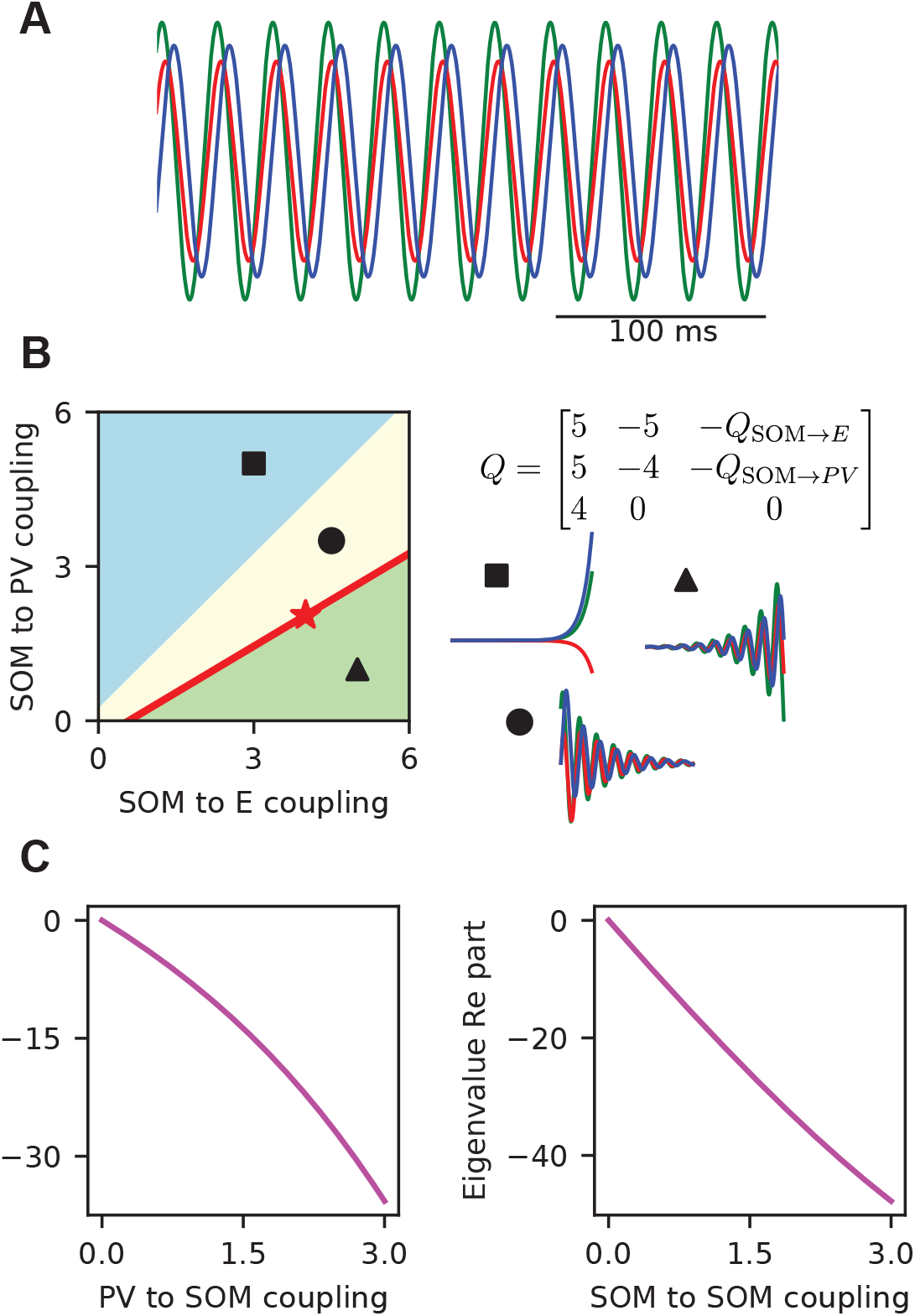
Linear mean-field analysis and phase diagram. (A) Linear mean-field dynamics at the parameter set indicated by the red star in (B). (B) Phase diagram of the linear mean-field system, with SOM→E strength on the x-axis and SOM→PV strength on the y-axis. Colors indicate different dynamical regimes: pale blue, coupling matrix with one positive real eigenvalue and a pair of complex conjugates with negative real parts; pale yellow, one negative real eigenvalue and a pair of complex conjugates with negative real parts; pale green, one negative real eigenvalue and a pair of complex conjugates with positive real parts. The red line marks the locus of purely imaginary eigenvalues, and the red star indicates the parameter point corresponding to (A). (C) Effect of adding additional inhibitory connections to the mean-field system at the red-star point. Left: SOM→SOM inhibition. Right: PV→SOM inhibition. In both cases, the system transitions from having a pair of purely imaginary eigenvalues to a pair of complex conjugates with negative real parts; moreover, increasing the strength of these added connections shifts the real part further into the negative range.

Next, we consider the system such that all *Q*_*∗*_s are fixed to the base linear model 10 values except *Q*_*SOM*→*E*_ and *Q*_*SOM*→*PV*_, which are treated as free parameters, and to investigate the dynamical regimes of the system. In order to characterize the dynamical regimes, the eigenvalues of the matrix

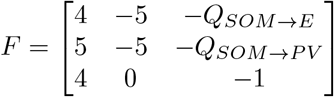

should be investigated. As shown in Appendix B, for any choice of *Q*_*SOM*→*E*_ and *Q*_*SOM*→*PV*_, the matrix *F* always possesses one real eigenvalue and a pair of complex conjugate eigenvalues.

Fig. 6B illustrates the dynamical regimes of

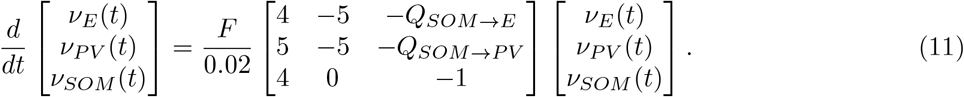

The red line, 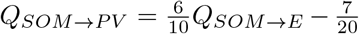, represents the dynamics where the solutions of the system are pure oscillatory. The point shown on the red line (by a red star) represents the setting mentioned above (where *Q*_*SOM*→*E*_ = 4 and *Q*_*SOM*→*PV*_ = 2.05). Below the red line (pale green), is the regime where the real part of the complex eigenvalues is positive and the real eigenvalue is negative and therefore the system is unstable (more precisely, the solutions are growing oscillations). Above the red line, the pale yellow region, is the regime where the real part of the complex eigenvalues is negative and the real eigenvalue is also negative. Therefore, the pale yellow regime is a stable regime or, more precisely, a damped oscillation. Pale blue is where the real part of complex eigenvalues is negative but the real eigenvalue is positive and consequently, the system is unstable. The boundary between pale blue and pale yellow is the line *h* = 0 or more precisely *Q*_*SOM*→*PV*_ = *Q*_*SOM*→*E*_ + 0.25 where the real eigenvalue is 0.

### Effects of PV→SOM and SOM→SOM couplings on the linear model

Here, we consider the base linear model 10 and firstly wonder what happens if we add a PV→SOM coupling to it. Hence, we need to investigate the system:

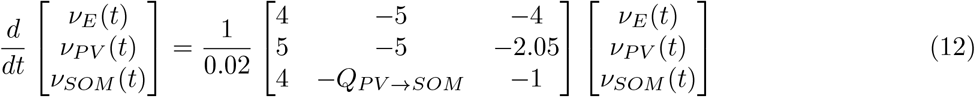

where *Q*_*PV* →*SOM*_ *>* 0 represents the PV to SOM coupling.

In order to investigate the effect of a non-zero *Q*_*PVtoSOM*_, eigenvalues of the matrix

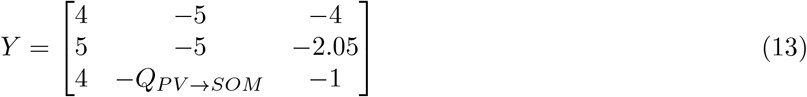

need to be analyzed. The determinant of Y is 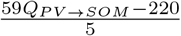, and needs to be equal to the multiplication of its eigenvalues. Therefore, if 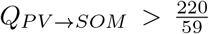, *Y* has at least one positive eigenvalue and so the solutions of equation 12 are asymptotically unstable. In view of this fact, we restrict our analysis to 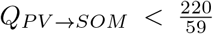. Through an analytical examination provided in Appendix C, it turns out that the solutions (for 0 *< Q*_*PVSOM*_ 220*/*59) are damped oscillations and as one increases *Q*_*PV→SOM*_, the oscillations exhibit stronger damping. The real part of the complex eigenvalues of 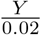 which represents the damping factor, can be seen as a function of *Q*_*PV* →*SOM*_ in Fig. 6C.

Secondly, we wonder what effect adding a SOM→SOM coupling can have on the base linear model 10.

To this aim, we need to investigate the eigenvalues of:

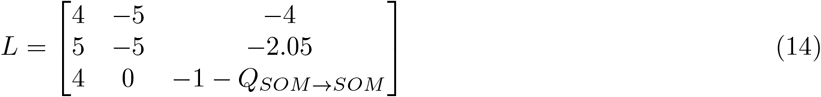

where *Q*_*SOM*→*SOM*_ *>* 0 represents the SOM to SOM coupling. By the examinations provided in Appendix D, it turns out that for any 0 *< Q*_*PV* →*SOM*_, the solutions of equation 14 are damping oscillations and as one increases *Q*_*SOMtoSOM*_, the oscillations exhibit stronger damping. The real part of the complex eigenvalues of 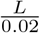 which represents the damping factor, can be seen as a function of *Q*_*SOM*→*SOM*_ in Fig. 6C.

## Discussion

In this work, we designed a mean-field model of E-PV-SOM cells that reproduces a large variety of experimental observations and provides some predictions. By employing linear stability analyses, we identified a robust mechanism of neural oscillation in E-PV-SOM networks, which emerges solely from the specific connectivity between SOM and PV interneurons. Remarkably, the structure of this connectivity matches anatomical observations of cell-type-specific synaptic organization.

We first designed a low-dimensional mean-field formulation of E-PV-SOM networks that accurately captures the population dynamics in the Canonical Microcircuit Network Oscillation (CAMINOS) model (Tahvili et al., 2025), recently introduced in the context of spiking networks with parameters calibrated from anatomical and electrophysiological data. We employed the master equation formalism developed in (Di Volo et al., 2019) and extended it to E-PV-SOM spiking neural networks with synaptic delay (employing the approach developed in (Tahvili and Destexhe, 2024)). Although this mean field model does not introduce a new methodology *per se*, the proposed E–PV–SOM mean-field formulation represents a substantial improvement over previous E/I models, as it can reproduce a wide range of experimental observations on the distinct roles of interneuron subtypes in shaping gamma oscillations and network stability. First, the mean-field model accurately reproduces the emergence of gamma oscillations at around 40 Hz, with SOM cells firing 6–7 ms later than excitatory and PV neurons, consistent with experimental observations (Onorato et al., 2020a). Second, the model captures key findings from optogenetic experiments in the primary visual cortex involving the suppression of SOM and PV neurons (Veit et al., 2017; Chen et al., 2017b). Importantly, our mean-field model, consistent with direct numerical simulations of the spiking network, demonstrates that PV interneurons primarily determine the oscillation frequency, while SOM interneurons play an essential role in shaping and sustaining the oscillatory amplitude. Furthermore, PV cells play a central role in maintaining network stability against hypersynchronous, epileptic-like dynamics. When PV activity is suppressed beyond a certain threshold (approximately 70 %), the oscillatory state in the meanfield model loses stability, leading to a bifurcation toward a high-activity regime of about 500 Hz, which represents the mean-field counterpart of a hypersynchronous epileptic state.

Our mean-field model also predicts that the frequency of oscillations in the E-PV-SOM model can cover alpha, beta and gamma ranges when varying properly the density ratio between PV and SOM cells in the network. Accordingly, the model also predicts that interregional differences in the ratio between PV and SOM cells could account for the dominant frequencies observed in cortical power spectra: alpha/beta oscillations prevailing in higher-order regions such as the prefrontal cortex, and faster gamma oscillations dominating in lower-order sensory areas such as the visual cortex (Vinck et al., 2025; Bastos et al., 2015; Vezoli et al., 2020, 2021; Murray et al., 2014; Honey et al., 2012; Buffalo et al., 2011; Vinck et al., 2023). Thanks to its capacity to reproduce the emergent dynamics of biologically realistic E–PV–SOM spiking networks, this mean-field model can serve as a building block for larger-scale frameworks that explicitly incorporate distinct inhibitory cell types.

Such a framework opens also new possibilities for modeling the emergence of multiple rhythms in interconnected E–PV–SOM modules, where different interneuron classes may dominate distinct frequency bands while remaining dynamically coupled. In addition, this mean-field formulation could be employed to investigate the organization of oscillatory power within a single cortical column across layers, particularly given the well-documented gradients in PV and SOM cell densities (Tremblay et al., 2016). Finally, since these gradients differ across species, our model could help predict interspecies variations in the population dynamics of cortical columns arising from distinct PV and SOM connectivity patterns and laminar distributions (Medalla et al., 2023).

To uncover the fundamental mechanism giving rise to oscillations in the E–PV–SOM model, we further simplified our mean-field formulation. We first removed the synaptic delay (*τ*_*D*_ = 0), since it is well known that *τ*_*D*_ *>* 0 represents a classical mechanism for generating gamma oscillations at a frequency proportional to 1*/τ*_*D*_ (Brunel and Wang, 2003). We also removed the spike-frequency adaptation time scale, thereby reducing our mean-field model to a three-dimensional system of ordinary differential equations (ODEs) with nonlinear transfer functions that depend on the biophysical parameters of the network and the distinct properties of each cell type.

Interestingly, this reduced mean field model still exhibits stable oscillations whose qualitative features closely match those of the original biologically realistic model. In particular, SOM cells always fire after E and PV neurons. Second, increasing either the PV → SOM or SOM → SOM coupling strength abolishes the oscillatory regime. These results demonstrate that the oscillations observed in our model do not rely on ad hoc temporal parameters such as synaptic timescales or adaptation, but rather constitute an intrinsic property of the three-population E–PV–SOM circuit. Interestingly, the emergence of oscillations depends critically on the specific structural organization of the network, characterized by weak PV → SOM and SOM→→ SOM coupling, as observed experimentally (Pfeffer et al., 2013).

We further extended our reductionist approach by considering a linear mean-field model. This approach isolates the essential connectivity motif among E, PV, and SOM populations. In this linear formulation, the only parameters are the coupling weights, that is, the connection strengths between distinct neuronal populations. An important advantage of this model is its analytical tractability, which allows for a formal investigation of the circuit’s dynamical properties. We showed that this linear system can sustain oscillations in the absence of SOM → SOM and PV → SOM connections, corresponding to the canonical architecture observed in anatomical studies. Moreover, the system becomes progressively less prone to oscillations as the strength of SOM→SOM and PV→SOM coupling increases. Interestingly, we analytically demonstrate that in this framework SOM cells always exhibit delayed activity with respect to E and PV neurons. This is a remarkable result, as it demonstrates that the oscillations observed in our biologically realistic model, and most likely those emerging in cortical circuits of E-PV-SOM cells, arise from a specific mechanism driven by the asymmetric connectivity between distinct interneuron types, namely SOM and PV cells.

Another interneuron subtype that has recently received attention is the vasoactive intestinal peptide-expressing (VIP) (Pfeffer et al., 2013; Veit et al., 2023). In this work, we focused on the development of E–PV–SOM circuits to elucidate the oscillatory mechanisms that this fundamental microcircuit can generate. Nevertheless, our mean-field approach—and the subsequent reduction to a linear analytical model—can readily be extended to include VIP interneurons. Although such an extension goes beyond the scope of the present study, investigating the influence of VIP neurons on gamma oscillations represents a promising future direction, well-suited to the methodological framework established here.

An important limitation of our study is the point-neuron approximation. Distinct interneuron types are known to have different dendritic targeting and integration properties; for example, SOM cells primarily project to the distal dendrites of excitatory pyramidal cells, whereas PV cells target their perisomatic regions (Tremblay et al., 2016). These morphological and integrative differences could be incorporated into our framework by introducing cell-type-specific synaptic delays or integration time constants. These differences may be important given the potential role of bursting excitatory neurons in generating gamma oscillations (Onorato et al., 2020b, 2025). However, to highlight the fundamental structural role of connectivity in generating oscillations, we deliberately chose to exclude such additional temporal parameters in the present work. Future studies could extend our model in this direction to assess how dendritic integration and spatial structure further shape and modulate oscillatory dynamics.

## Methods

### Spiking Network

To model the dynamics of the neuronal membrane potential *V* (*t*), we employ the adaptive exponential integrate-and-fire (AdEx) model (Brette and Gerstner, 2005). The governing equations are given by

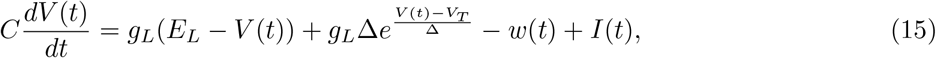

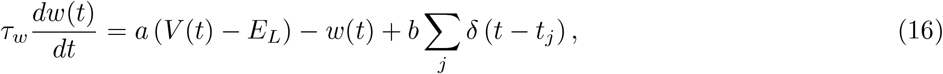

where *C* denotes the membrane capacitance, *g*_*L*_ the leak conductance, *E*_*L*_ the resting potential, *V*_*T*_ the effective threshold, and Δ the slope factor controlling the sharpness of spike initiation. The current *I*(*t*) corresponds to the sum of all synaptic inputs. The variable *w*(*t*) represents the adaptation current, which contributes a negative feedback to the membrane voltage. Each time a spike occurs at *t*_*j*_, *w* is increased by *b* due to the integration of the Dirac delta function *δ*(*t* − *t*_*j*_). The parameter *a* encodes the subthreshold adaptation factor, *τ*_*w*_ defines the adaptation time constant, and the spike threshold was set to *V*_*T*_ + 5Δ. After spiking, the membrane potential is reset to *V*_reset_ and held constant for a refractory period *T*_ref_.

Parameter values were chosen to reflect electrophysiological properties of excitatory pyramidal neurons, parvalbumin-expressing (PV) interneurons, and somatostatin-expressing (SOM) interneurons (McCormick et al., 1985; Kawaguchi and Kubota, 1996; McGarry et al., 2010; Tremblay et al., 2016). Notably, PV cells lack spike-frequency adaptation, whereas both excitatory and SOM cells exhibit it. Moreover, spike-frequency adaptation tends to be more pronounced in SOM cells compared to pyramidal regular-spiking cells (Romero-Sosa et al., 2021; Yavorska and Wehr, 2016; Erisir et al., 1999). The leak potential *E*_*L*_ was tuned across cell types to capture average firing rates reported in V1, with higher activity in PV cells and lower rates in SOM and pyramidal cells (Ma et al., 2010). All other parameters were set to biologically realistic values within standard ranges. A complete list of parameters is provided in Table 1.

**Table 1:** Neurons’ parameters.

| Parameter | E, PV, SOM |
| --- | --- |
| $V_T$ | -50 mV |
| $\Delta$ | 2, 0.5, 2 mV |
| $T_{\text{ref}}$ | 5 ms |
| $\tau_w$ | 500 ms |
| $a$ | 4, 0, 4 nS |
| $b$ | 50, 0, 80 pA |
| $C$ | 200 pF |
| $g_L$ | 10 nS |
| $E_L$ | -65, -65, -55 mV |
| $V_{\text{reset}}$ | -65 mV |

Synaptic interactions were modeled as conductance-based inputs of the form

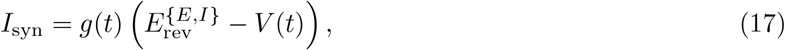

where *V* (*t*) is the postsynaptic potential and 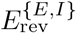 is the reversal potential of excitatory (E) or inhibitory synapses. The synaptic conductance *g*(*t*) evolves according to

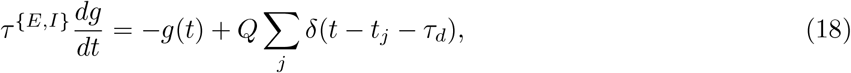

where *τ* ^*{E,I}*^ is the excitatory (E) or inhibitory (I) synaptic decay time constant, *τ*_*d*_ the synaptic delay, *δ*(*t* − *t*_*j*_ − *τ*_*d*_) representing a presynaptic spike arriving at the delayed time, and *Q* the quantal conductance (denoted *Q*_*X*→*Y*_ for connections from population *X* to *Y*). Synaptic strengths were set based on experimental estimates (Destexhe et al., 1998). A complete list of synaptic parameters is provided in Table 2.

**Table 2:** Synaptic parameters.

| Parameter | Value |
| --- | --- |
| $\tau_d$ | 2 ms |
| $\tau^E$ | 2 ms |
| $\tau^I$ | 8 ms |
| $E_{\text{rev}}^E$ | 0 mV |
| $E_{\text{rev}}^I$ | -80 mV |
| $Q_{\{\text{PV}, \text{SOM}\} \rightarrow \{\text{E}, \text{PV}\}}$ | 5 nS |
| $Q_{\text{E} \rightarrow \text{E}}$ | 2 nS |
| $Q_{\text{E} \rightarrow \{\text{PV}, \text{SOM}\}}$ | 3 nS |
| $Q_{\text{Ext} \rightarrow \{\text{E}, \text{PV}\}}$ | 2.5 nS |

The network consisted of *N*_*E*_ = 8000 excitatory pyramidal neurons and *N*_*I*_ = 1500 inhibitory interneurons, equally divided into PV and SOM subpopulations (*N*_*PV*_ = 750, *N*_*SOM*_ = 750). Connectivity was random and sparse: any presynaptic neuron formed a synapse onto a postsynaptic target with probability *p* = 0.07, consistent with experimental observations (Campagnola et al., 2022). In the reference model (Fig. 1A), two connections are absent in line with experimental evidence: PV→SOM and SOM→SOM (Pfeffer et al., 2013). Thus, apart from the analyses of Section 5, we set *Q*_*PV* →*SOM*_ = 0 and *Q*_*SOM*→*SOM*_ = 0.

External drive was modeled as independent Poisson spike trains at 4 Hz, generated by *p* · *N*_*E*_ excitatory neurons, each providing inputs with quantal conductance *Q*_Ext_ = 2.5 nS to excitatory and PV cells.

Population activity for each cell class was calculated as

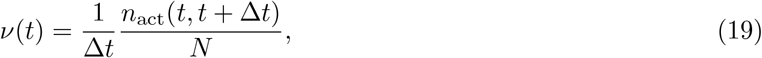

where *N* is the size of the population and *n*_act_(*t, t* + Δ*t*) the number of spikes in the interval (*t, t* + Δ*t*). Unless otherwise stated, we used Δ*t* = 0.1 ms. Resulting activity traces *ν*(*t*) were convolved with a Gaussian kernel of width 2 ms for visualization.

Simulations were implemented in Python using the Brian2 package (Stimberg et al., 2019). Differential equations were integrated using the Euler method with a timestep of *dt* = 0.1 *ms*.

### Transfer function derivation for the mean-field

For the mean-field description, we derived transfer functions ℱ _*E*_, *ℱ* _*P*_ *ℱ* _*V*_, _*SOM*_ that map presynaptic firing rates onto the output rate of each neuron type. The approach follows the shot-noise diffusion approximation (Fourcaud and Brunel, 2002; Zerlaut et al., 2016; Di Volo et al., 2019), in which the effective input statistics are estimated and then converted into an output rate through a fluctuation-dependent threshold.

#### Input statistics

Given presynaptic rates *ν*_*E*_, *ν*_*PV*_, *ν*_*SOM*_, *ν*_ext_, the corresponding event rates are

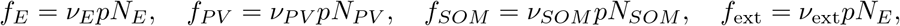

with connection probability *p* and population sizes *N*_*X*_. Each input contributes a mean synaptic conductance *µG*_*X*_ = *Q*_*X*_*τ*_*X*_*f*_*X*_. The total conductance is

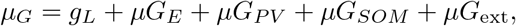

yielding an effective time constant *T*_*m*_ = *C*_*m*_*/µ*_*G*_. The mean subthreshold potential is approximated as

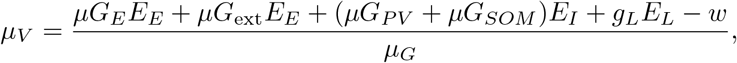

with *w* = 0 for PV cells.

#### Voltage fluctuations

The postsynaptic impact of a single event of type *X* is

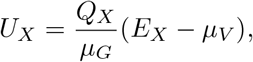

where *E*_*X*_ is the corresponding reversal potential. The variance of membrane potential fluctuations is

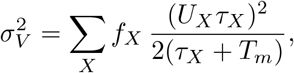

and the effective correlation time is

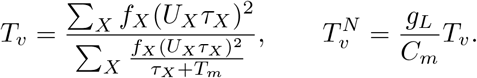

#### Effective threshold and transfer function

To capture nonlinear dependence on input statistics, the effective threshold is expressed as a second-order polynomial in the normalized variables (*µ*_*V*_ − *µ*_*V* 0_)*/Dµ*_*V* 0_, (*σ*_*V*_ − *σ*_*V* 0_)*/Dσ*_0_, and 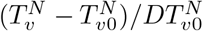, with coefficients *P*_0_, …, *P*_9_ fitted from AdEx single-cell simulations. All numerical parameters and fitted coefficients are reported in Table 3 and Table 4. The output firing rate is then

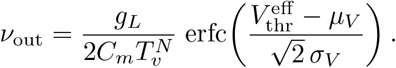

**Table 3:**
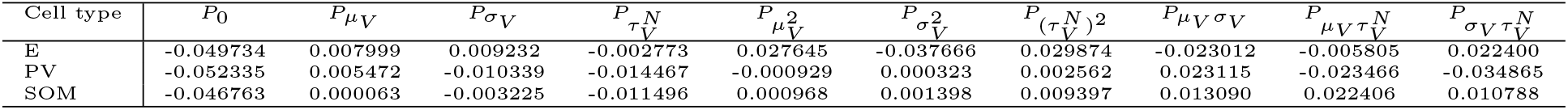
Fitted parameters for each cell type.

| Cell type | $P_0$ | $P_{\mu_V}$ | $P_{\sigma_V}$ | $P_{\tau_V^N}$ | $P_{\mu_V^2}$ | $P_{\sigma_V^2}$ | $P_{(\tau_V^N)^2}$ | $P_{\mu_V \sigma_V}$ | $P_{\mu_V \tau_V^N}$ | $P_{\sigma_V \tau_V^N}$ |
| --- | --- | --- | --- | --- | --- | --- | --- | --- | --- | --- |
| E | -0.049734 | 0.007999 | 0.009232 | -0.002773 | 0.027645 | -0.037666 | 0.029874 | -0.023012 | -0.005805 | 0.022400 |
| PV | -0.052335 | 0.005472 | -0.010339 | -0.014467 | -0.000929 | 0.000323 | 0.002562 | 0.023115 | -0.023466 | -0.034865 |
| SOM | -0.046763 | 0.000063 | -0.003225 | -0.011496 | 0.000968 | 0.001398 | 0.009397 | 0.013090 | 0.022406 | 0.010788 |

**Table 4:** Baseline fit parameters for each cell type.

| Cell Type | $\mu_{V0}$ | $D\mu_{V0}$ | $s_{V0}$ | $Ds_{V0}$ | $T_{vN0}$ | $DT_{vN0}$ |
| --- | --- | --- | --- | --- | --- | --- |
| E | -0.06 | 0.05 | 0.004 | 0.006 | 0.5 | 1 |
| PV | -0.06 | 0.01 | 0.004 | 0.006 | 0.5 | 1 |
| SOM | -0.05 | 0.01 | 0.004 | 0.006 | 0.5 | 1 |

## Appendix A

### Proof 1.

*Suppose that P is a strictly positive diagonal* 3*∗* 3 *matrix like* diag(*a*1, *a*2, *a*3) *and M* = *PAP* ^−1^. *The eigenvalues of M are the same as those of A and if x is an eigenvector of A then Px is an eigenvector of M*. *The eigenvector of M corresponding to the eigenvalue α* + *iω is a*_*1*_ *x*_*1*_ *a*_*2*_ *x*_*2*_ *a*_*3*_ *x*_*3*_ ^*T*^. *Since a_i_ are strictly positive numbers, the phase of a*_*i*_*x*_*i*_ *is the same as the phase of x*_*i*_ *and therefore it is enough to prove for M*.

*We take P* = diag(1, *Q*_*PVtoE*_, *Q*_*SOMtoE*_) *and therefore*

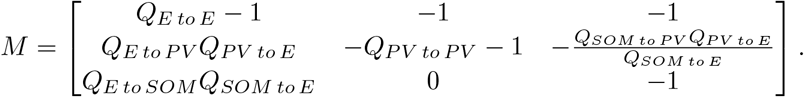

*Without loss of generality, suppose that the eigenvector of α* + *iω is ν*_1_ *ν* 1 *and, therefore*,

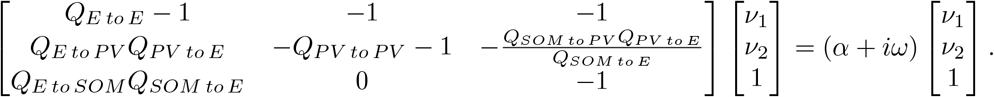

*From the third row*, 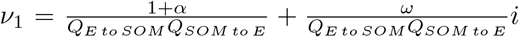 *is derived. Since, ω, Q*_*EtoSOM*_ *and Q*_*SOMtoE*_ *are larger than zero, we have* 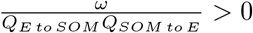 *and therefore E activity has a phase advance to the SOM activity*.

*From the first row, we have* (*Q*_*EtoE*_ − 1)*ν*_1_ − *ν*_2_ − 1 = (*α* + *iω*)*ν*_1_ *and it follows that ν*_2_ = ((*Q*_*EtoE*_ − 1) − *α* − *iω*)*ν*_1_ − 1. *By substituting* 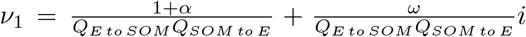, *we conclude that the imaginary part of ν*_2_ *is*

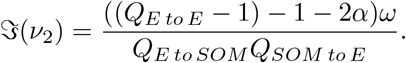

*Now it remains to prove that this imaginary part is strictly positive. To do this, it is enough to prove that Q*_*EtoE*_−1 *>* 1+2*α. Suppose, for the sake of contradiction, that K* = (−(*Q*_*EtoE*_−1)+1+2*α*) ≥ 0. *We already knew that the matrix A, consequently M, has three eigenvalues α*+*iω, α*−*iω and r. It is known that the sum of eigenvalues is equal to the sum of diagonal entries and therefore r* = (*Q*_*EtoE*_ −1)−(*Q*_*PVtoPV*_ +1)−(1+2*α*). *On the other hand, since r is an eigenvalue of M then* det(*M* − *rI*) = 0 *and it follows that*

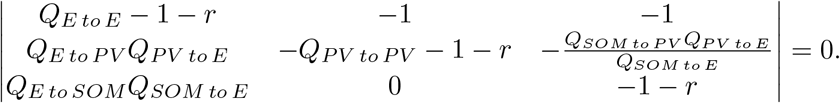

*Expanding the determinant along its third row and taking K* = (−(*Q*_*EtoE*_ − 1) + 1 + 2*α*) *results in* det(*M* −*rI*) = *Q*_*PV to E*_*Q*_*SOM to PV*_ *Q*_*E to SOM*_ +*Q*_*E to SOM*_ *Q*_*SOM to E*_*K* +(*K* +*Q*_*PV to PV*_)(*Q*_*E to PV*_ *Q*_*PV to E*_ + *K*(*Q*_*PVtoPV*_ + 2 + 2*α*). *On the other hand from Q*_*EtoE*_ *>* 0 *and K* ≥ 0 *we have* 2 + 2*α >* 0. det(*M* − *rI*) *is a sum of positive terms and some of these terms are strictly positive (like Q*_*PVtoE*_*Q*_*SOMtoPV*_ *Q*_*EtoSOM*_, *Q*_*PVtoPV*_ *Q*_*EtoPV*_ *Q*_*PVtoE*_, *etc*.*) and therefore* det(*M* − *rI*) *>* 0 *which is a contradiction. Therefore, we proved that Q*_*EtoE*_ − 1 *>* 1 + 2*α and it follows that ℑ*(*ν*_2_) *>* 0.

## Appendix B

We show that for any choice of *Q*_*SOM*→*E*_ and *Q*_*SOM*→*PV*_, the matrix *F* possesses one real eigenvalue and a pair of complex conjugate eigenvalues. Eigenvalues of *F* are the roots of its characteristic polynomial and therefore are the solutions of *λ*^3^ + 2*λ*^2^ + (6 + 4*Q*_*SOM*→*E*_)*λ* + (20*Q*_*SOM*→*E*_ − 20*Q*_*SOM*→*PV*_ + 5) = 0. Let us denote 6 + 4*Q*_*SOM*→*E*_ and 20*Q*_*SOM*→*E*_ − 20*Q*_*SOM*→*PV*_ + 5 by *g* and *h*, respectively. The cubic discriminant of the polynomial is then Δ = −4*g*^3^ + 4*g*^2^ + (36*g* − 32)*h* − 27*h*^2^. View Δ as a quadratic in *h*: Δ(*h*) = −27*h*^2^ + (36*g* − 32)*h* + (4*g*^3^ − 4*g*^2^). The discriminant of Δ(*h*) (as a quadratic in *h*) is *D*_*h*_ = − 16(3*g* 4)^3^. We know that *g* = 6 + 4*Q*_*SOM*→*E*_ and therefore *g* is always larger than 6 resulting in *D*_*h*_ *<* 0. Since the coefficient of *h*^2^ in Δ(*h*) is −27 *<* 0, the quadratic Δ(*h*) opens downward and having negative discriminant - never meets the *h* axis. Thus Δ(*h*) *<* 0, for all *h* and so we proved that the cubic discriminant of *F* is always smaller than 0. Δ *<* 0 dictates that the characteristic polynomial of *F* has exactly one real and two non-real conjugate roots. Hence, we showed that no matter what *Q*_*SOM*→*E*_ and *Q*_*SOM*→*PV*_ are, *F* has exactly one real eigenvalue and a pair of complex conjugate eigenvalues.

## Appendix C

Let us denote *Q*_*PVtoSOM*_ by *x* and so the characteristic polynomial of *Y* is 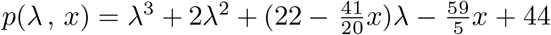. By reasoning like the one provided for matrix *F* (by analyzing the discriminant of the polynomial), we conclude the matrix *Y* has a pair of complex conjugate eigenvalues and a real eigenvalue. *p*(−2, *x*) = −7.7*x* and 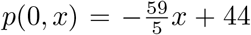 and therefore for any 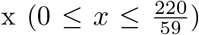, we have *p*(−2, *x*) ≤ 0 and *p*(0, *x*) ≥ 0. Hence, by the intermediate value theorem, it turns out that the real eigenvalue of *Y* is in [− 2, 0]. Let us denote the real eigenvalue of *Y* by *r*(*x*). We have *p*(*r*(*x*), *x*) = 0 and by differentiating both side we derive

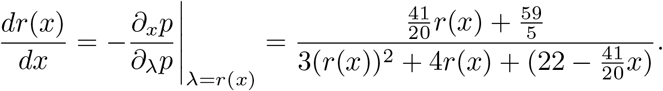

We already know that 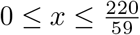 and 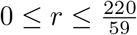 and therefore both the numerator and denominator are positive and therefore *r*(*x*) is a strictly increasing function of *x*. On the other hand, the trace of *Y* equals the sum of its eigenvalues and so 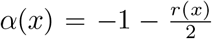 where *α*(*x*) is the real part of the complex conjugate eigenvalues. Consequently, it follows that *α*(*x*) is negative and a strictly decreasing function of *x*.

## Appendix D

Let us denote *Q*_*SOMtoSOM*_ by *x* and therefore the characteristic polynomial of *L* is *p*(*λ, x*) = *λ*^3^ + (*x* + 2)*λ*^2^ + (*x* + 22)*λ* + (5*x* + 44). By reasoning using the discriminant of the polynomial similar to the one provided before one can show that the matrix *L* has a pair of complex conjugate eigenvalues and a real eigenvalue for any *x* ≥ 0. Let’s show the eigenvalues by *α*(*x*) *± β*(*x*) and *r*(*x*). Implicit differentiation of p(r(x), x) = 0 gives

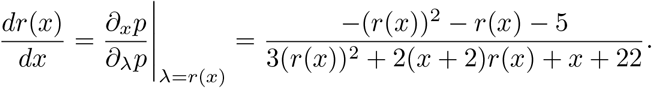

For any *r*(*x*), the numerator is always negative. On the other the denominator is

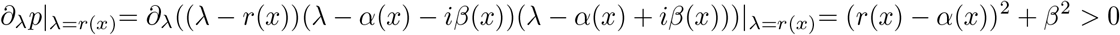

and it follows that r’(x) is always negative. And so r(x) is a negative decreasing function of *x*, starting from r(0) = -2.

Equality of the summation of eigenvalues and the trace of *L*, results in 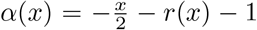. By similar analysis (omitted for brevity), one can show that *α*(*x*) starts from *a*(0) = 0, decreases strictly to its global minimum around −1.3 at around *x*^*∗*^ = 6 and then it strictly increases to its limit, 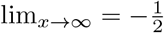.

## Acknowledgements

FT and MdV were supported by the French Ministry of Higher Education (Ministére de l’Enseignement Supérieur) and the project LABEX CORTEX (ANR-11-LABX-0042) of Université Claude Bernard Lyon 1 operated by the ANR. MV was supported by an ERC starting grant (850861) SPATEMP, DFG VI Grants (908/5-1 and 908/7-1; 505660261; 520285844; SPP LOOPS), an NWO VIDI Grant, and the Dutch Brain Interface Initiative (DBI2).

